# Disentangling leaf-microbiome interactions in *Arabidopsis thaliana* by network mapping

**DOI:** 10.1101/2022.04.05.487248

**Authors:** Kaihang Li, Kexin Cheng, Haochen Wang, Qi Zhang, Yan Yang, Yi Jin, Xiaoqing He, Rongling Wu

## Abstract

The leaf microbiota plays a key role in plant development, but a detailed mechanism of microbe-plant relationships remains elusive. Many genome-wide association studies (GWAS) have begun to map leaf microbes, but few has systematically characterized the genetics of how microbes act and interact. Previously, we integrated behavioral ecology and game theory to define four types of microbial interactions – mutualism, antagonism, aggression, and altruism, in a microbial community assembly. Here, we apply network mapping to identify specific plant genes that mediate the topological architecture of microbial networks. Analyzing leaf microbiome data from an Arabidopsis GWAS, we identify several heritable hub microbes for leaf microbial communities and detect 140-728 SNPs responsible for emergent properties of microbial network. We reconstruct Bayesian genetic networks from which to identify 22-43 hub genes found to code molecular pathways related to leaf growth, abiotic stress responses, disease resistance and nutrition uptake. A further path analysis visualizes how genetic variants of Arabidopsis affect its fecundity through the internal workings of the leaf microbiome. We find that microbial networks and their genetic control vary along spatiotemporal gradients. Our study provides a new avenue to reveal the “endophenotype” role of microbial networks in linking genotype to end-point phenotypes in plants. Our integrative theory model provides a powerful tool to understand the mechanistic basis of structural-functional relationships within the leaf microbiome and supports the need for future research on plant breeding and synthetic microbial consortia with a specific function.

**IMPORTANCE:** It is found that plant genes act as microbiome gatekeepers to select which microbes get to live inside the leaves for health. Many genome-wide association studies (GWAS) have begun to map leaf microbes, but few has systematically characterized the genetics of how microbes act and interact. This work illustrates a more comprehensive picture of the genetic architecture underlying the leaf microbiome by network mapping. This study also dissects how genetic variants affect its fecundity by direct path and indirect path through microbial network, revealing the “endophenotype” role of microbial networks in linking genotype to end-point phenotypes. Future studies could benefit from this work to improve understanding the underlying genetic mechanisms that govern the relationships between plants and their microbiomes, and to manipulate plant genetic system to reconfigure microbiome. Plants could become more efficient at selecting their microbial partners to improve their health, resilience, and productivity.

## Introduction

Plants, as the main producer and sustainer of terrestrial ecosystems, are colonized by a variety of microorganisms that form complex microbiomes, including bacteria, fungi, archaea and protists (1, 2). These microorganisms interact dynamically with plants and influence their hosts’ growth and development. Plant microbiome can enhance plants growth directly or indirectly through increasing abiotic stresses tolerance (3–5), nutrient acquisition (6, 7), disease resistance (8, 9), and pathogen inhibition via the synthesis and excretion of antibiotics (10). The composition and functioning of the microbiomes even can predict plant health (11) and help mitigate the negative consequences of climate change (12).

Most studies on plant microbiomes have focused on rhizosphere microbial communities and their functioning rather than on those of phyllosphere. The phyllosphere microbiomes may play essential but often overlooked roles in nutrient acquisition, abiotic stress tolerance, and disease suppression of plants (13). Experimental studies have demonstrated that the phyllosphere harbors diverse microbial communities that influenced ecosystem functioning (14–19). Recent studies stress the importance of understanding the mechanisms underlying how plant-microbe interactions in the phyllosphere could influence host survival and fitness in the context of global change (20, 21). Several studies have illustrated that host genotypes can influence the composition of the leaf microbiome (22–24), but little is clear how the plant shapes its leaf microbiota and how the leaf microbiome contributes to plant phenotypic traits (25–27).

It has been recognized that microbial interactions affect plant traits, plant evolution (28, 29) and ecosystem function (30). Despite tremendous efforts to reveal the molecular mechanisms of microbial interactions (31, 32), their role in modulating plant function through the context of ecological networks has been little studied. This may be due to the fact that the microbiota is a highly-packed ecosystem, making it extremely difficult to discern and quantify individual microbial interactions. By integrating behavioral ecology and game theory, Wu and team developed mathematical descriptors for quantifying and characterizing different types of microbial interactions, including mutualism (two microbes promotes each other), antagonism (two microbes inhibit each other), aggression (a stronger microbe is aggressive to a weaker microbe), and altruism (one microbe benefit the other), in ecological communities at any large scale (33, 34). These mathematical descriptors, named Wu’s descriptors for convenient mention, have been biologically validated by designing and conducting a series of cultural experiments using the fish and bacteria (33, 35). Based on Wu’s descriptors, S. Wu et al. (36) have further formulated conceptual hypotheses on the strategic choice of organisms’ behavior in complex communities. X. He et al. (37) introduced Wu’s descriptors into a GWAS setting, proposing a network mapping tool to study the genetic architecture of microbial networks.

Here, we implement network mapping to dissect the internal workings of the leaf microbiome for *Arabidopsis*, using the published GWAS data from a well-designed multi-trial and multi-year experiment (24). In this study, key plant genetic variants that influenced leaf hub microbes responsible for *A. thaliana* fitness were identified. Through our reanalysis, we attempt to illustrate a more comprehensive picture of the genetic architecture underlying the leaf microbiome of *A. thaliana*. First, network mapping allows us to reconstruct four ecologically different types of microbial networks based on Wu’s descriptors. Thus, we can map host genes for the topological architecture of microbe-microbe interactions and interdependence. Second, it has been increasingly clear that microbial traits are not only determined by key host QTLs, but also through their epistatic networks (37). Network mapping can draw the host genetic landscape of microbial interactions. Taken together, network mapping can discern both hub microbes in the leaf microbiome and hub QTLs in the genetic architecture of the host and characterize the impact of these different types of hubs on plant fitness. We further implement path analysis to chart the roadmap of genotype-phenotype linking through microbial interactions as the endophenotypes.

## RESULTS

### Ecological networks of the leaf microbiota

Fig.1 illustrates the mutualism, antagonism, aggression and altruism interaction networks reconstructed with Wu’s descriptors for the leaf microbiome of Arabidopsis at four different sites in two years. In the mutualism network, we calculate the relative abundances of the secondary leaders over the primary leaders, the tertiary leaders over the secondary leaders and the secondary leaders over the followers in the eight experiments (in green, Fig. 1). All these relative values are obviously larger than 0.618, which complies with the golden threshold hypothesis, stating that a larger microbe tends to cooperate with a smaller microbe when the relative size of the latter to the former is beyond 0.618 (36). By comparing the abundance of hawks and doves in the aggression network, we find that the ratios of the hawks-doves to the hawks ranged from 0.373 - 0.798 (the upper tier) and the ratios of the doves to the hawks-doves is between 0.557 - 0.765 (the lower tier) (in yellow, Fig. 1). Many of these ratios are lower than 0.618, which is consistent with the second aspect of the golden threshold hypothesis, stating that a larger microbe tends to exploit a small microbe when the relative size of the latter to the former is below 0.618 (36). In the altruism network, the relative abundance of egoists over altruists at both the upper and lower tiers are obviously larger than 0.382 (in purple, Fig.1). This is in agreement with the Fibonacci retracement mark hypothesis, stating that for its self-interest, a smaller microbe may “trick” a larger microbe when the relative size of the former to the latter is beyond 0.382 (36). In the antagonism network, the relative abundance of smaller antagonists over larger antagonists ranges from 0.668 to 0.903 (in red, Fig.1), which is in agreement with the surrender-resistance hypothesis. These hypotheses regarding microbial interactions can unravel the quantitative mechanisms of how microbes cope with others to gain their maximum benefits and, ultimately, affect community assembly structure, organization, and function. It seems that both bacteria and fungi on the Arabidopsis leaves empirically obey these hypotheses (Fig.1, Supplemental Table 2).

**Fig. 1.**
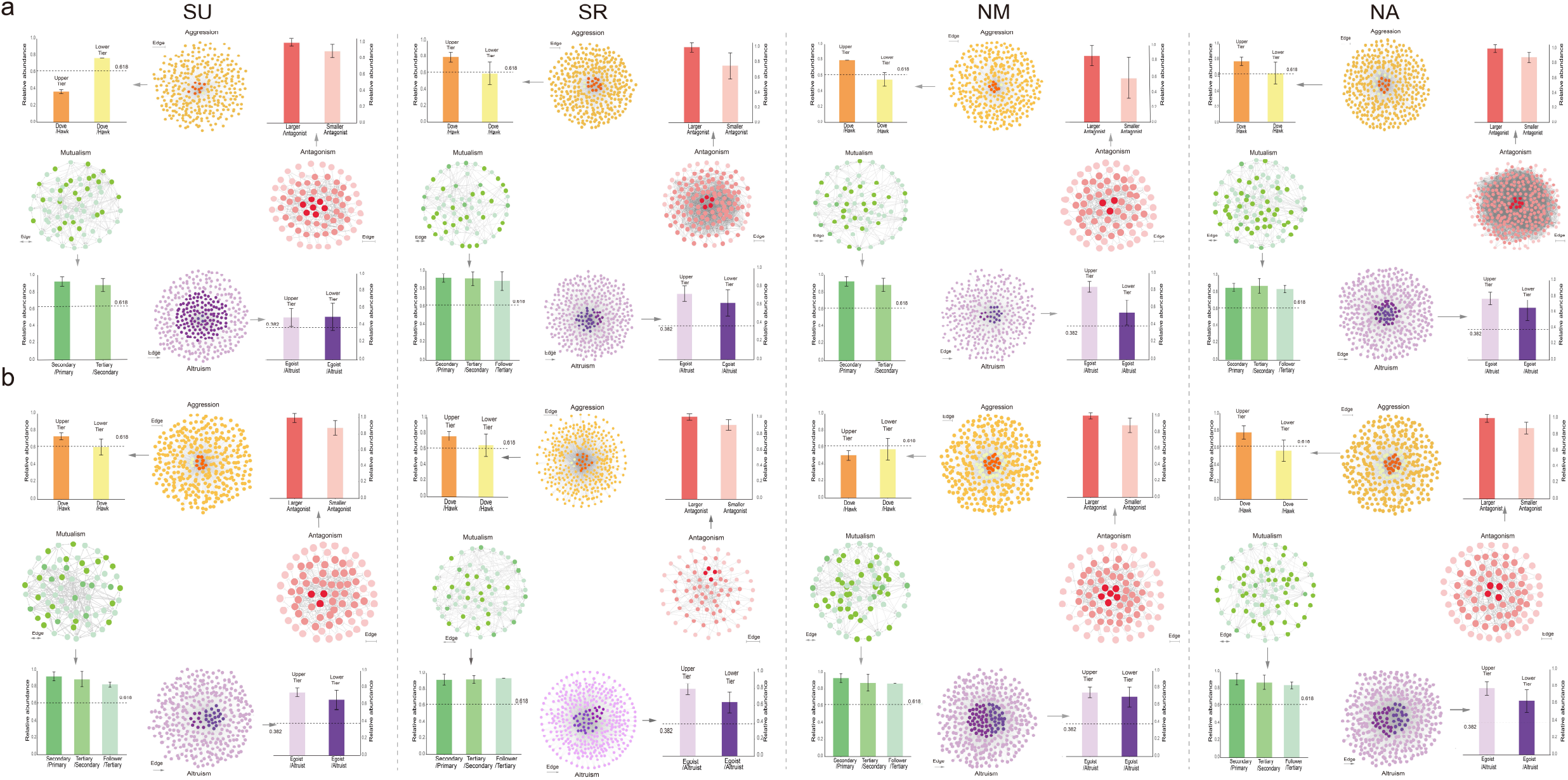
Social networks of microbiome in the leaf of *Arabidopsis thaliana*. Left: Mutualism network constructed by doubly-arrowed edges representing microbes cooperation. Right: Antagonism network composed of doubly-T-shaped edges denoting the mutual conflict of the microbes. Top: Aggression network represented by singly-T-shaped edges specifying how a microbe (as a hawk) aggresses upon others (doves) (color metrics indicate three hierarchies of aggression). Bottom: Altruism network characterized by singly-arrowed edges which illustrates how a microbe (altruist) benefits other microbes (egoists) at its own expense (color metrics indicate three hierarchies of altruism).

### Hub microbes and OTU heritability

We identify 96 hub OTUs in microbial networks from eight experiments (Fig.2, Supplemental Table 3). These hubs are from bacterial phyla: Proteobacteria (46 OTUs), Bacteroidetes (5 OTUs), Actinobacteria (4 OTUs), and Firmicutes (2 OTUs), and fugal phyla: Basidiomycota (19 OTUs), Ascomycota (18 OTUs), and unclassified fungi (2 OTUs). OTU1 (Proteobacteria, *Sphingomonas* sp. TSBY-34) is the only one detected in each site and year (Fig. 3). OTU2 (Proteobacteria, *Pseudomonas* sp.) and OTU 213 (Ascomycota, *Tetracladium* sp.) are detected in 7 experiments, while fungi OTU 201 (Basidiomycota, *Itersonilia perplexans)* is detected in 6 experiments. OTU 11 (Proteobacteria, uncultured), OTU 5 (Proteobacteria, *Variovorax* sp.), OTU 202 (Ascomycota, *Tetracladium maxilliforme*) and OTU 206 (Basidiomycota, *Tremellales* sp.) are detected in 5 experiments.

**Fig. 2.**
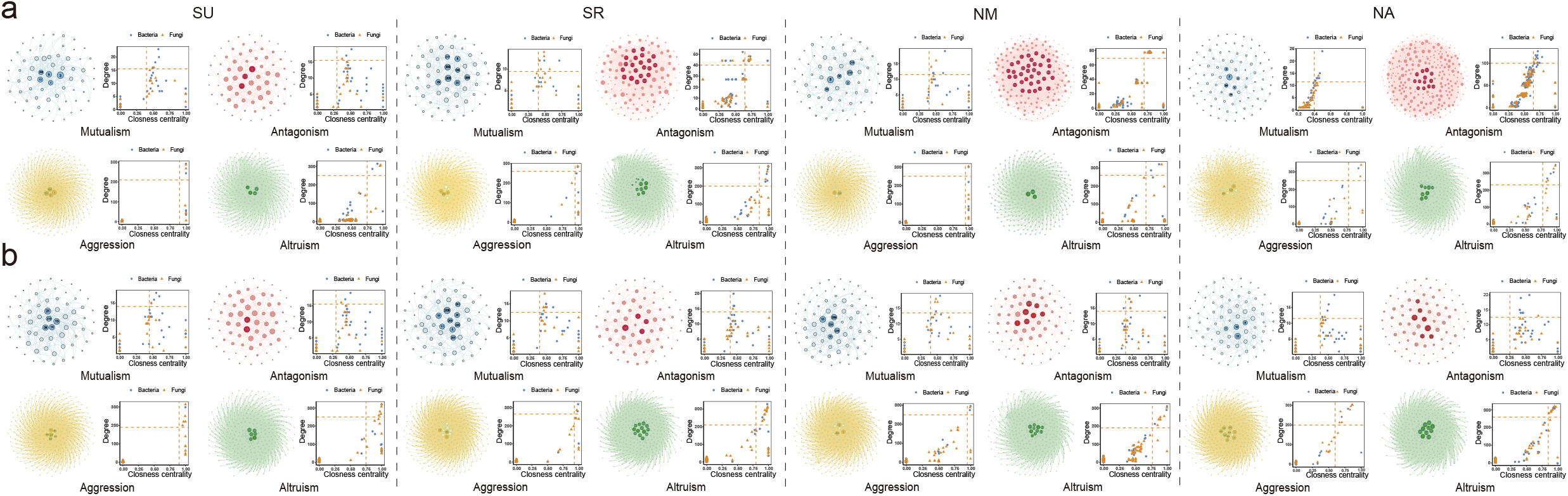
Ecologically hub OTUs of the co-occurrence network. In each network, hub microbes are highlighted in border colors. The distribution of ‘Hub microbes’ in four different microbial networks was based on degree and closeness centrality values. These two values of each OTU within each network were given at the right. The red dotted line represents the screening cutoffs of ‘Hub microbes’ corresponding to each network. Visualization was done with Gephi for four microbial networks.

**Fig. 3.**
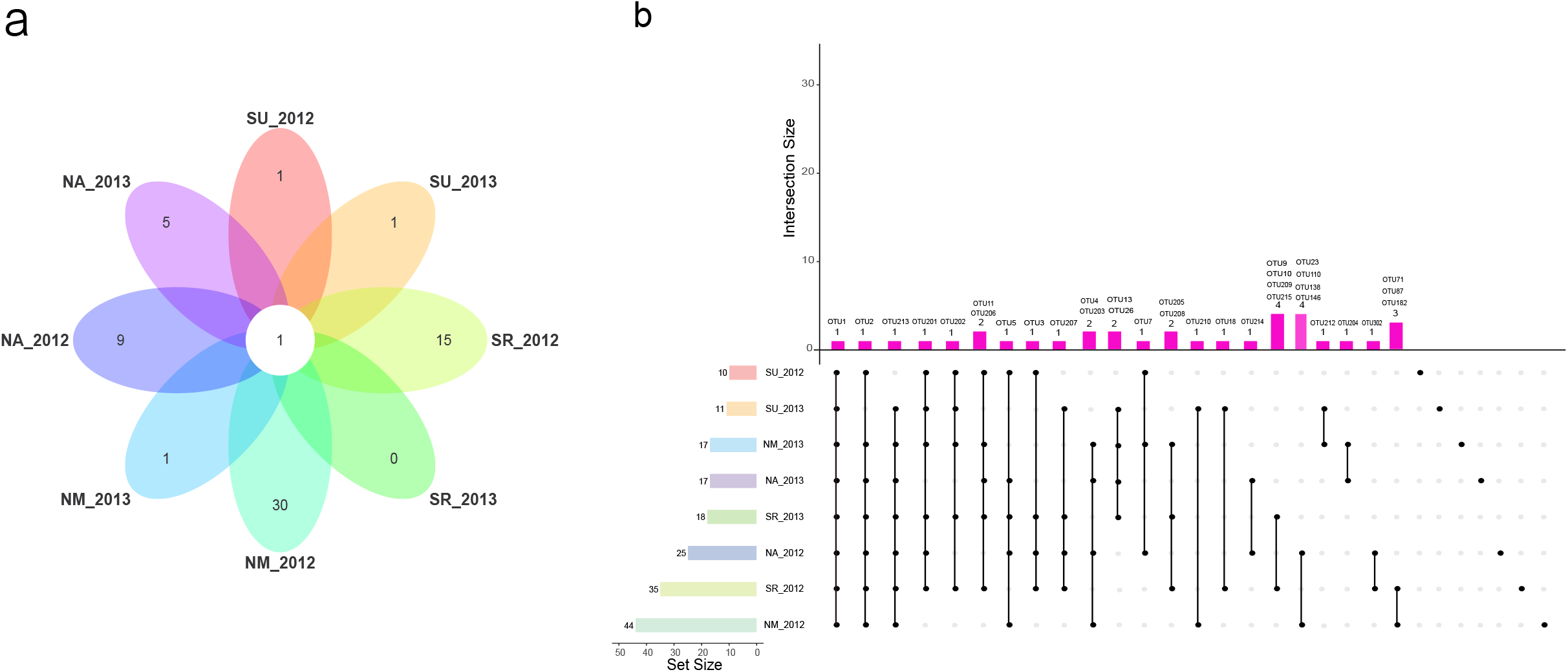
Venn plot for hub OTUs. **(A)** Flower plot shows the number of common and unique hub OTUs in 8 experiments, (**B)** UpSet plot for 8 experiments. Horizontal bars on the left represent the total number of hub OTUs in each experiment. Deep pink bars represent the number of hub OTUs of each intersection indicated by connected dots.

We calculate the broad-sense heritability (*H^2^*) of the abundance of individual OTUs in 8 experiments using the function *lmer* in *lmer4* R package (24) (Supplemental Table 4). We did not find large *H^2^* for the OTUs, ranging from 0 to 0.2287. Across all eight experiments, 15 hub OTUs are detected to be heritable (with *H^2^* > 0.10).

### Hub QTLs for microbial networks

We calculate six network property indices to describe networks’ emergent properties (Supplemental Fig. 1) and map significant QTLs for each index in 8 experiments (Supplemental Fig. 2). The population structure (*Q*) and relative kinship (*K*) are performed by Admixture and EMMAX, respectively, to control spurious associations (38). Supplemental Fig.3 illustrates the quantile-quantile plots of the P value distributions based on the model without consider *Q* and *K*, the *Q* model, and the *Q* + *K* model. We calculate the genomic inflation factor λ of the three models in R (Supplemental Table 5). The model with λclose to 1 is chosen to perform association analysis. We also investigate candidate genes within ~10 kb windows on each side of associated SNPs by software PLINK. The genes of R^2^ > 0.8 were retained. Finally, we identify 139-727 significant SNPs responsible for microbial network properties in 8 experiments (MAF>0.05) (Supplemental Table 6). We find that SNP-based heritability (h^2^) for network properties varies from 0 to 12.79 % (Supplemental Table 7).

In the Bayesian QTL networks (Fig. 4, Supplemental Table 8), 28 - 66 hub QTLs are excavated in 8 experiments. There are 40 pleiotropic QTLs including those in the region of *ADA2B (AT4G16420*) and *AT5G02880 (HAL3A*), detected to influence multiple properties of various microbial networks (Supplemental Table 8). Hub gene *AT1G78070* is identified in two experiments (NM_2012 and SR_2013), while *AT1G23060 (MDP40*) identified in NA_2012 and SU_2013. Gene annotation analysis suggests that a number of hub genes detected are biologically relevant, playing roles in leaf growth, abiotic stress responses, disease resistance, and nutrition uptake (Supplemental Table 8).

**Fig. 4.**
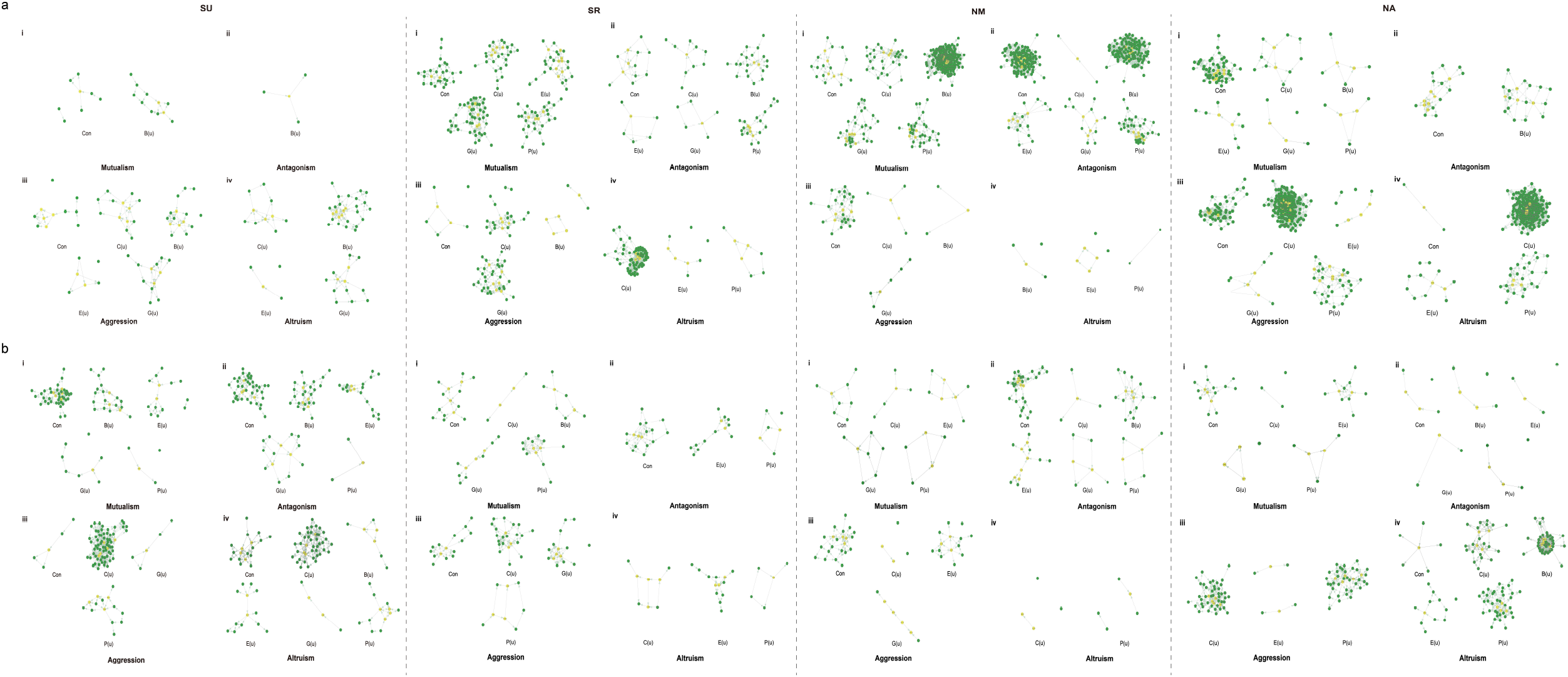
Bayesian networks of the significant SNPs from microbial networks. **(A)** mutualism, **(B)** antagonism, **(C)** aggression and **(D)** altruism. Each node reports a SNP and hub QTLs (SNPs) were colored in green. The network characteristic indices were described by connectivity (Con), closeness (C(u)), betweenness (B(u)), eccentricity (E(u)), eigencentrality (G(u)), and PageRank (P(u)).

Hub gene *ALG10* (*AT5G02410*) encodes alpha1,2-glucosyltransferase (ALG10) that is required for oligosaccharide biosynthesis and subsequently for normal leaf development and abiotic stress response (39). The inactivation of ALG10 in Arabidopsis results in the activation of the unfolded protein response and increased salt sensitivity. Hub gene *AT5G11250* encodes an atypical TIR-NBS protein acting as a regulator of the hormonal response to stress and is required for plant survival and robustness to environmental perturbations (40). Gene *DDF2* encodes a member of the DREB subfamily A-1 of ERF/AP2 transcription factor family (DDF2), which is expressed in all tissues but most abundantly expressed in rosette leaves and stems. Overexpression of this gene results in the reduction of gibberellic acid biosynthesis and helps the plants increase their tolerance to high-salinity levels. Hub gene *ADA2B* (*AT4G16420*) encodes a transcriptional co-activator ADA2b in Arabidopsis responses to abiotic stress. It is required for the expression of genes involved in abiotic stress either through modulation of histone acetylation in the case of salt stress or affecting nucleosome occupancy in low temperatures response (41).

Hub gene *GLIP2* (*AT1G53940*), encoding a GDSL motif lipase/hydrolase–like protein, plays a role in pathogen defense via negative regulation of auxin signaling (42). AtLPK1 is a plasma membrane-localized L-type lectin-like protein kinase 1, which is encoded by hub gene *AT4G02410* in this study. Overexpression of AtLPK1 confers the pathogen resistance to infection by *Botrytis cinerea* and regulates salinity response in *Arabidopsis thaliana*, which implicates that AtLPK1 plays essential roles at both abiotic and biotic stress response (43). Hub gene *AT1G25550*, encoding nitrate-inducible NIGT1.1/HHO3 proteins, involves in regulating nitrate signaling and phosphorus starvation signals in Arabidopsis (44). Hub gene *HAL3A* (*AT3G18030*) expresses HAL3-like protein A which is related to salt and osmotic tolerance and plant growth (45).

### Microbial networks as endophenotypes for linking host genotype to phenotype

We further implement path analysis to dissect the role of microbial networks in linking host genotype (at significant SNPs) to end-point phenotype-fecundity. Although all SNPs display a sizable direct effect on fecundity, they also affect fecundity through the indirect effects of microbial networks as endophenotypes (Fig. 5). For example, SNP7, residing in the genomic region of gene *AT1G12570*, is the ortholog of maize IPE1 gene which is involved in pollen exine development. The IPE1 mutant exhibits defective pollen exine and is male sterile (46). *AT1G12570* positively affects fecundity in a direct way, but it also affects fecundity through positive indirect effects of betweenness and eigenvector in the aggression and altruism network in the experiment of SR_2013.

**Fig. 5.**
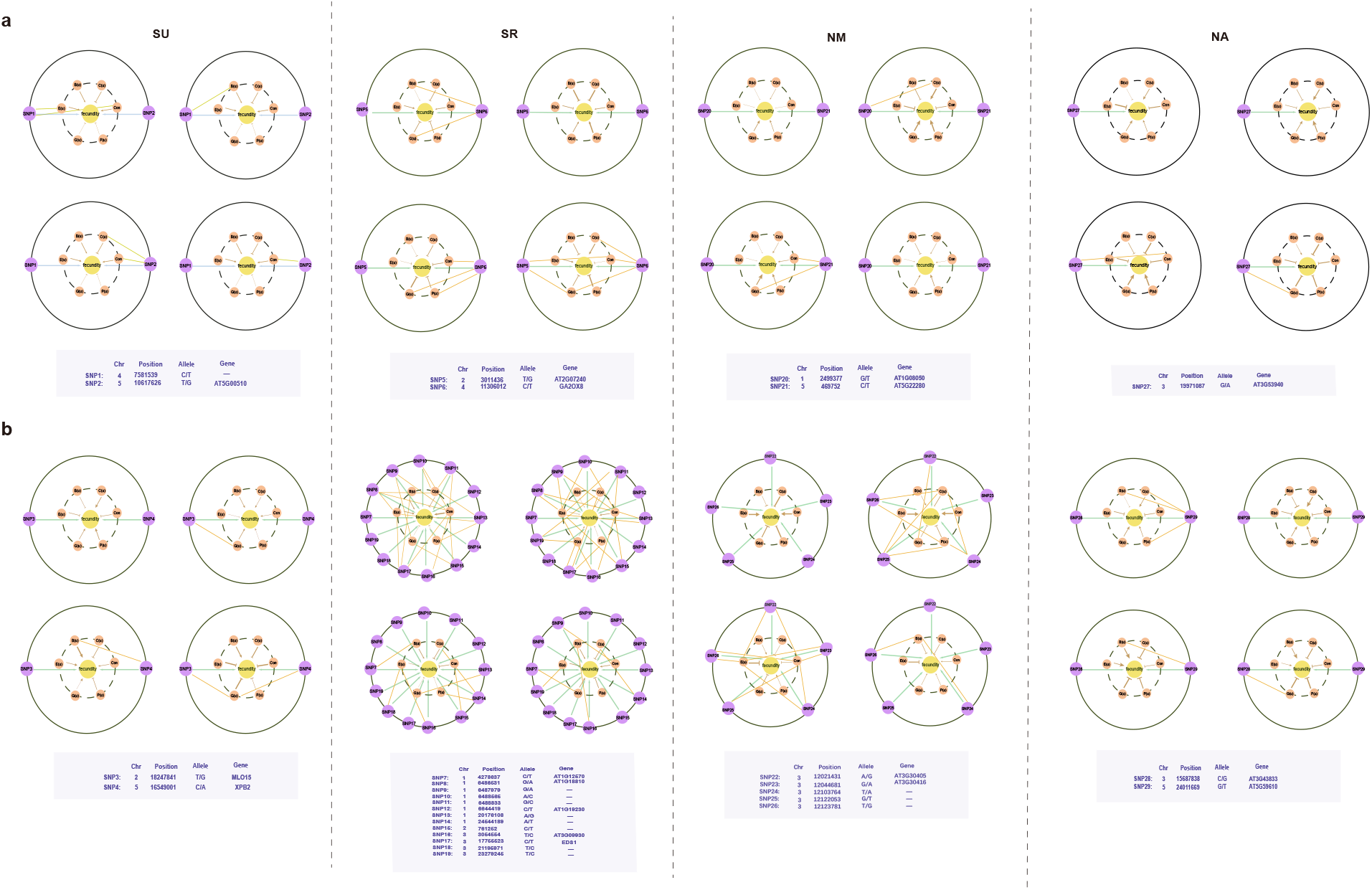
Path analysis revealing how QTLs (outer) affect fecundity as a final phenotype (inner) through microbial networks as an “endophenotype” (middle) (described by differences of six emergent property indices). Path coefficients are denoted by directed lines from SNPs to fecundity (green) and from networks to fecundity (brown). Arrowed line and T-shaped line represent a positive and negative impact of path, respectively. Correlation coefficients between SNPs and networks are denoted by blue lines. The thickness represents the magnitude of path and correlation coefficients.

## Discussion

As many studies have been devoted to understanding the diversity of the rhizosphere microbiota, increasing attention has been paid to studying the phyllosphere, despite its disconnection to soil, often considered to be a site for relatively few microbes (47). In this study, we apply an advanced network mapping theory for mapping the genetic architecture of microbial interaction networks in the leaf microbiome of Arabidopsis (24). From this analysis, we gain new insight into how microbial interactions mediate the leaf microbiome assembly and how plant genes affect plant traits

The most remarkable feature of this network mapping study is founded on Wu’s descriptors, a group of mathematical equations to characterize and discern different types of ecological interactions, namely, mutualism, antagonism, aggression and altruism, which occur in highly dense microbial community assembly (33–35). Each of these interaction types describes a different aspect of network topology and function, which can better explain how each microbe interacts with every other microbe and how microbial cooperation and competition are reciprocally shifted in response to environmental change (36).

Plants are able to regulate a few beneficial microbial taxa to maximize their fitness (48). Identifying hub taxa is one of the advantages for network analysis (49). Hubs are highly connected in a microbiome, which are responsible for microbiome structure and maintain the dynamics of the community (50). Some hub species could act as a mediator to curate a healthier community indirectly even though themselves do not promote plant growth(51). Our network mapping using Arabidopsis phyllosphere reveals that phyla Proteobacteria, Actinobacteria, Bacteroidetes, and Firmicutes are dominant groups. Individual strains of Proteobacteria (Sphingomonas, Rhizobium) and Actinobacteria (Microbacterium, Rhodococcus) as keystone species play an important role in affecting community structure (52). These two phyla are also considered as ecological hub OTUs in this study; especially *Sphingomonas* sp. TSBY-34 serves as a keystone species in all 8 experiments. We calculate broad sense heritability for individual OTU abundances and identified 15 OTUs that are both heritable and hubs in at least one of the eight experiments.

In a large-scale longitudinal field study of the maize rhizosphere microbiome, heritable taxa identified were diverse, including 26 Alphaproteobacteria, 9 Betaproteobacteria, 12 Actinobacteria, 6 Verrucomicrobia, and 8 Bacteroidetes (53). The use of synthetic communities (SynComs) allows for dissecting how one or few community members affect *A. thaliana* and how host genes affect microbiome composition (22, 54, 55). Even one single bacterial genus (Variovorax) has an ability to maintain root growth in a complex microbiome (56). A simplified SynComs is able to rescue plant from root rot disease (57). Heritable hubs may serve as the components in the SynComs system in the future research on plant–microbe interactions.

B. Brachi et al. (24)identified host genotype effects on the relative abundance of microbial hubs and LSP across sites and years. Their analysis detected a few significant genes for the abundance of heritable hubs. Our network mapping characterizes previously undetected QTLs that mediate microbial interactions. There are 40 pleiotropic hub genes, including *ADA2B* (*AT4G16420*) and *AT5G02880* (*HAL3A*), which are detected to influence multiple types of microbial networks and in 8 experiments. Gene annotation suggests that a number of hub genes detected are biologically relevant, playing roles in leaf growth, abiotic stress responses, disease resistance and nutrition uptake. Some of these genes are also found to affect root development (Supplemental Table 8).

Much research has shown that plant genes is a key driver to maintain the balance of leaf microbiomes. Plants impaired in genetic networks embrace a dysbiosis leaf microbiome in structure and composition (14). The plant genetic network links the leaf microbial community to plant health. For example, a study found that the immunity and cell surface component structuring genes *A. thaliana* mutant harbored less diverse bacterial community in the leaf endosphere relative to the wild type and induced leaf chlorosis (15). Our understanding of how plant genotypes impact colonization of specific microorganisms will be instrumental in spurring next-generation plant breeding strategies (58, 59).

By integrating network mapping and path analysis, we can characterize how SNPs determine plant phenotype through their direct effects or the indirect effect through leaf microbial networks. We identify these two different paths for SNP-fecundity links. Indirect genetic effects related to these SNPs are mediated through multiple microbial interaction types.

In the near future, modulating the balance of the leaf microbial community by regulating host genetic networks may become a novel approach to improve crop traits and maintain sustainable agricultural development. Our network mapping could hold a great promise to achieve this goal.

## Conclusion

In this study, we quantify the networks of various interaction types for the leaf microbiome in Arabidopsis thaliana. We dissect leaf-microbiome interactions by network mapping to reveal the genetic architecture of microbial interactions and identified hub plant genes. We conduct path analysis to dissect the roadmap from each of these SNPs to fecundity into the direct path and the indirect paths through microbial network, revealing the “endophenotype” role of microbial networks in linking genotype to end-point phenotypes.

## Materials and methods

### Leaf microbiome experiments

B. Brachi et al. (24) performed a GWAS for the leaf microbiome in *A. thaliana*. The study included a panel of 198 *A. thaliana* accessions planted with 2 replicates in the spring of 2012 and 2013 at four sites located in Ullstorp (lat: 56.067, long: 13.945; SU in short), Ratchkegården (lat: 55.906, long: 14.260; SR in short), Ramsta (lat:62.85, long 18.193; NM in short), and Ådal (lat: 62.862, long 18.331; NA in short). The bacterial and fungal abundance of each leaf sample was measured by16s/ITS rRNA gene sequencing. Accessions were genotyped by a high-throughput sequencing technique, obtaining 186,161 SNPs after quality control. For a detailed description of experimental design, sampling strategy, microbial sequencing, and SNP genotyping, refer to B. Brachi et al. (24).

In this study, we choose 200 bacterial OTUs and 200 fungal OTUs at the top-abundance from each site for microbial network inference. These numbers of OUTs account for the top 95.89% of bacterial relative abundance and top 95.50% of fungal relative abundance, respectively, (Supplemental Table 1). OTU1-200 are listed as bacteria and OTU200-400 as fungi. B. Brachi et al. (24) measured fecundity for each plant, whose genetic architecture is disssected using both SNPs and microbes.

### Quantitative microbial networks

Using Wu’s descriptors, we reconstruct and visualized 400-node interaction networks based on mutualism, antagonism, aggression, and altruism, for eight year-site experiments. OTU abundance is chosen as the trait that determines the strategies of microbial interactions. We calculate the relative OTU abundance of primary, secondary leaders, tertiary leaders and followers in the mutualism network, the relative OTU abundance of two antagonists in the antagonism network, the relative OTU abundance of hawks and doves in the aggression network, and the relative OTU abundance of altruist and egoists in the altruism network (36). Different members were shown by the metric of colors.

Hub taxa are identified from each type of microbial network using the R *igraph* package. We calculate the degree of each node in a network. To reduce the bias, we statistically identify the hub taxa with the highest degree and closeness centrality (13, 37). Meanwhile, six network indices, including connectivity (Con), closeness (C(u)), betweenness(B(u)), eccentricity (E(u)), eigencentrality (G(u)), and Pagerank (P(u)) are calculated, as described previously (35).

### Mapping QTL networks underlying microbial networks

We apply a likelihood approach for detecting significant SNPs that are associated with each of the six network properties for eight year-site experiments (60). We further use *bnlearn* R package to reconstruct Bayesian genetic networks among the significant SNPs detected. SNP-SNP interactions are visualized by the *plot* function. Hub QTLs, playing a key role in genetic networks, are identified.

### Path analysis

We implement path analysis to test whether a significant SNP affects plant fecundity directly or through an indirect pathway of microbial interactions. Let *g* denotes the genotype, *y* denotes the network property, and *z* denotes the fecundity. We calculated the Pearson correlation between *y* (continuous) and *z* (continuous) across individuals, denoted as *r_yz_*. Let *r_gy_* and *r_gz_* denote the correlations between *g* and *y* and between *g* and *z*, respectively. We calculate the path coefficients *P_z←g_* from the equation

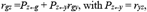

where *P_z←g_* is the direct path from g to *z* and *P_z←y_r_gy_* is the indirect path through microbial networks.

### Availability of data and materials

The data that support the findings of this study can be downloaded at http://dx.doi.org/10.1101/181198. The computer code can be freely downloaded at https://github.com/lenahe2006/leaf-microbiome-interactions for any purpose of research. All questions in data and computation can be addressed to the corresponding author.

## Acknowledgements

We thank Dr. Bergelson for supplying their leaf Arabidopsis microbiome and SNP data to us.

## Author’s information

Affiliations

**College of Biological Sciences and Technology, Beijing Forestry University, Beijing, People’s Republic of China**

Kaihang Li, Kexin Cheng, Haochen Wang, Qi Zhang, Yan Yang, Yi Jin, Xiaoqing He

**Center for Statistical Genetics, The Pennsylvania State University, Hershey, PA 17033 USA**

Rongling Wu

## Contributions

KL analyzed the data, and prepared figures and tables; KC analyzed the data, and prepared figures and tables; HW analyzed the data, and prepared figures and tables; QZ analyzed the data; YY analyzed the data; YJ revised the paper; XH conceived and wrote the paper; RW conceived and revised the paper; All authors authored or reviewed drafts of the paper and approved the final draft.

## Corresponding authors

Correspondence to Xiaoqing He or Rongling Wu

## Ethics declarations

### Ethics approval and consent to participate

Not applicable.

### Consent for publication

Not applicable.

## Supplemental information

### Supplemental Tables

**Supplemental Table 1** The relative abundance of top 200 OTUs of bacteria and fungi in each experiment.

**Supplemental Table 2** Social networks of leaf microbiome in each experiment.

**Supplemental Table 3** List of ecologically hub OTUs in all experiments.

**Supplemental Table 4** List of heritable hubs.

**Supplemental Table 5** The inflation factors of three models.

**Supplemental Table 6** The list of significant SNPs.

**Supplemental Table 7** SNP-based heritability estimates.

**Supplemental Table 8** The hub SNPs and gene description.

### Supplemental Figures

**Supplemental Fig.1** Heatmap of six emergent property indices. **(A)** The results of 4 experiments in 2012, **(B)** The results of 4 experiments in 2013. (a) Mutualism (b) Antagonism (c)Aggression (d) Altruism.

**Supplemental Fig.2** Manhattan plots of GWAS results for *Arabidopsis thaliana* genomic regions associated with emergent properties of leaf microbial networks including mutualism, antagonism, aggression and altruism in 8 experiments. In each circle from inside to outside, it represents the association results of six network indices respectively, connectivity (Con), closeness (C(u)), betweenness (B(u)), eccentricity (E(u)), eigenvector(G(u)) and pagerank (P(u)). The red dotted lines were reported significant threshold (-log10(P)≥5). Each dot in the plot represented an SNP, and significant SNPs were colored in red. **(A)** The results of 4 experiments in 2012, **(B)** The results of 4 experiments in 2013. (a) Mutualism (b) Antagonism (c)Aggression (d) Altruism.

**Supplemental Fig.3** Quantile-quantile plots of the P value distributions from three models in 8 experiments. **(A)** QQ plots of 4 experiments in 2012, **(B)** QQ plots of 4 experiments in 2013.

